# curatedPCaData: Integration of clinical, genomic, and signature features in a curated and harmonized prostate cancer data resource

**DOI:** 10.1101/2023.01.17.524403

**Authors:** Teemu D Laajala, Varsha Sreekanth, Alex Soupir, Jordan Creed, Federico CF Calboli, Kalaimathy Singaravelu, Michael Orman, Christelle Colin-Leitzinger, Travis Gerke, Brooke L. Fidley, Svitlana Tyekucheva, James C Costello

## Abstract

Genomic and transcriptomic data have been generated across a wide range of prostate cancer (PCa) study cohorts. These data can be used to better characterize the molecular features associated with clinical outcomes and to test hypotheses across multiple, independent patient cohorts. In addition, derived features, such as estimates of cell composition, risk scores, and androgen receptor (AR) scores, can be used to develop novel hypotheses leveraging existing multi-omic datasets. The full potential of such data is yet to be realized as independent datasets exist in different repositories, have been processed using different pipelines, and derived and clinical features are often not provided or unstandardized. Here, we present the *curatedPCaData* R package, a harmonized data resource representing >2900 primary tumor, >200 normal tissue, and >500 metastatic PCa samples across 19 datasets processed using standardized pipelines with updated gene annotations. We show that meta-analysis across harmonized studies has great potential for robust and clinically meaningful insights. *curatedPCaData* is an open and accessible community resource with code made available for reproducibility.

## INTRODUCTION

Prostate cancer is the most common cancer type amongst men with an estimated incidence of 268,490 new cases per year in the United States, with an estimated 34,500 deaths per year^1^. Molecular profiling of prostate cancer has led to insights into the relationship of genomic alterations and disease initiation, progression, and treatment response. However, no significant differences in disease free survival were found for patients that were stratified according to the 8-group prostate cancer (PCa) taxonomy defined by The Cancer Genome Atlas (TCGA) using single gene molecular alterations^2^. Additionally, when primary tumors were compared to metastatic tumor samples, few changes in the frequency of these genomic alterations were observed^2–4^.

A reliable molecular biomarker that stratifies aggressive vs. indolent disease is increased frequency of Copy Number Alterations (CNAs)^4–7^; however, this finding provides little mechanistic or therapeutically actionable insight. Recent studies have shown that combinations of alterations, namely *TP53 & RB1*^*8*^ and *CHD1 & MAP3K7*^*9*^, drive aggressive disease, suggesting that molecular subtyping in PCa is complex. Many efforts have been put forward to develop predictive gene expression signatures with the goal of identifying which patients will progress to lethal disease^10–16^. Some of these signatures have been clinically successful^11,17,18^; however, an overwhelming amount of gene expression profiling results lack replicability between studies resulting in inconsistent lists of candidate genes associated with PCa prognosis^19^. Additional challenges in reproducible PCa research remain. For example, the use of high-dimensional molecular data is dependent on thorough validation of the statistical models in diverse datasets. Similar concerns apply to molecular subtyping. Many of these challenges can at least partially be addressed by harmonization of the omic-data preprocessing and annotations, matched with manual curation of the clinicopathologic features and outcomes for easy application of multi-study statistical learning^20^ and cross-study validation^21^.

Data wrangling and data harmonization are critical for consistent, reproducible and benchmarked analysis of multi-omic cancer datasets. Efforts have been completed for ovarian cancer in the curatedOvarianData R package^22^, breast cancer in the curatedBreastData R package^23^, and across cancer types in the curatedTCGAdata R package^24^. These packages have advanced the field in many ways. To this end, the R user community has put great effort into developing R class objects that help end-users to utilize data across different types - such as transcriptomics, copy number alterations, and somatic mutations - and between studies that vary in their specific study characteristics. The *MultiAssayExperiment*-class^25^ (MAE) aggregates data of various types utilizing such R classes as *matrix, RaggedExperiment, SummarizedExperiment* across these data levels. This data class supports linking and simultaneous storage of sample or patient-level clinical metadata fields that can be easily processed and stored together with their corresponding ‘omics’ data.

In addition to the primary ‘omic’ data types themselves, such as gene expression measurements by RNA sequencing or microarrays, there are now an array of innovative approaches to develop molecular signatures and deconvolution methods to estimate cell types present in bulk tissue. The *immunedeconv*-package^26^ has proven to be a popular choice as a wrapper R package providing harmonized access to multiple popular cell type deconvolution methods such as EPIC^27^, ESTIMATE^28^, MCP-counter^29^, quanTIseq^30^, and xCell^31^. Estimating prevalences of different cell types in the tumor specimen has allowed for investigating the relationship between immune cell and other cell frequencies in a tumor sample with clinical outcomes^26–34^.

Given the value to the PCa research field in having a unified resource of molecular features across independent studies, we developed a curated, comprehensive, and harmonized PCa resource that contains multi-omic and clinical data from 19 PCa studies. The ‘omic’ data types were preprocessed and annotated, and clinical variables were mapped to the common data dictionary to ensure consistent annotation of the samples. Furthermore, we precomputed several prostate-specific genomic scores using the uniform preprocessed and annotated gene expression data sets. Namely, we conveniently provide Decipher^35^, Oncotype DX^36^, and Prolaris^37^ risk scores as well as Androgen Receptor (AR) scores^2^. These precomputed variables can be easily included in the downstream analyses as correlatives or phenotypic variables. Leveraging the MAE class, we supply the data in the *curatedPCaData* R package. The package provides open and accessible data and analysis pipelines with maximum flexibility for data analysts and prostate cancer researchers. We discuss the integrated datasets within the package and insights that have been gained by bringing together >3500 prostate tissue, primary PCa, and metastatic PCa tumor samples in one location: https://github.com/Syksy/curatedPCaData.

## RESULTS

The *curatedPCaData* package was developed using standardized workflows for raw data processing where available, mapping all clinical information for each dataset to a common data dictionary (**Table S1**), and ensuring gene symbols are consistent and up-to-date using HUGO Gene Nomenclature Committee (HGNC) symbols across all datasets and data types (**Figure S1**). To harmonize, organize, and manage all datasets and data types, the *curatedPCaData* package was built using the data structures for multi-omic data integration as implemented in the *MultiAssayExperiment* R package^25^. A summary of the key study characteristics of the 19 datasets contained in the *curatedPCaData* package are in **Table 1**.

**Table 1:**
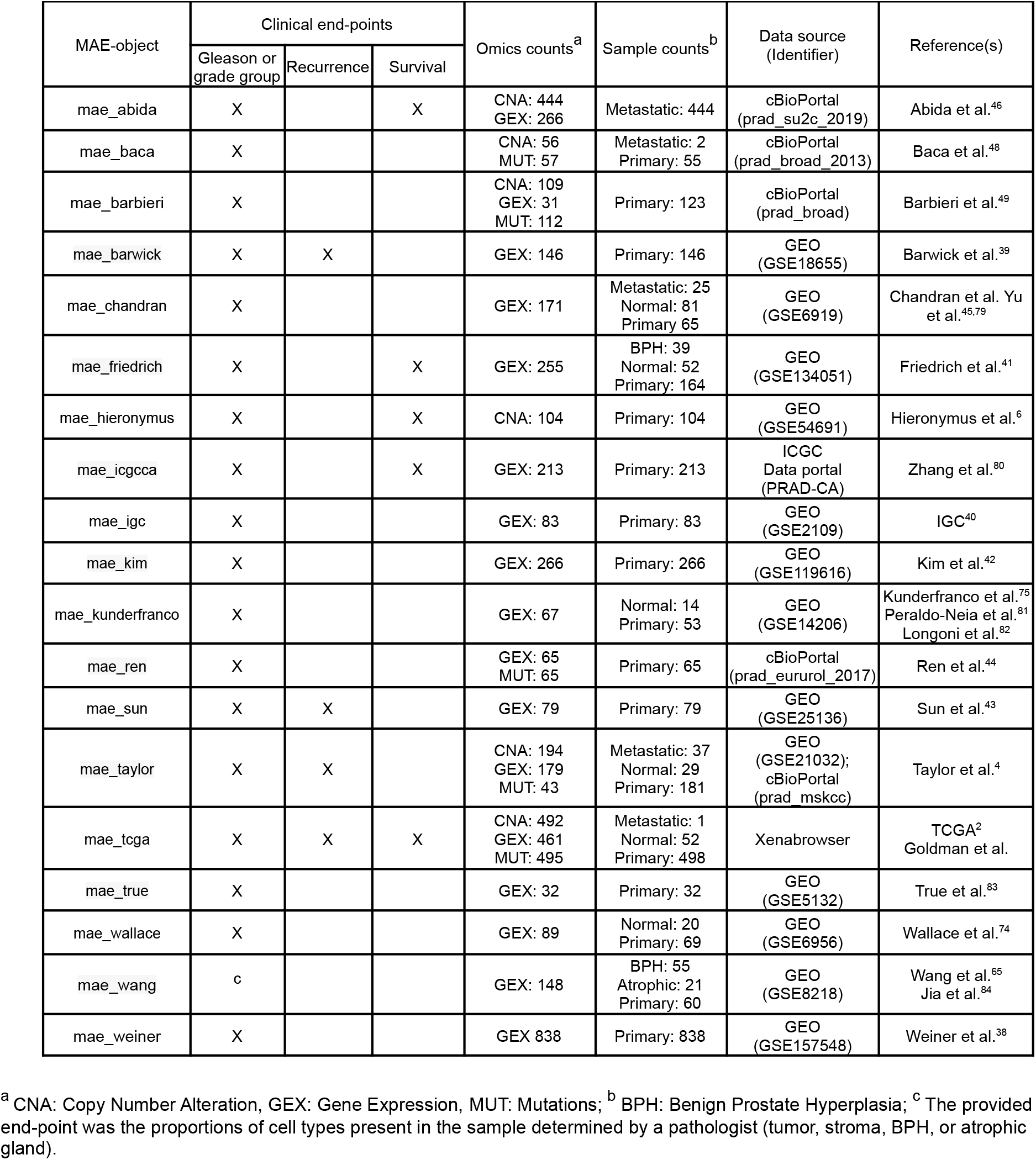
Summary of studies in *curatedPCaData* and their corresponding MultiAssayExperiment (MAE) object contents.

For reproducibility and to provide users with example code, all analyses and results presented in the following sections are made available as vignettes through the *curatedPCaData* package (**Table S2**).

### Molecular measurements are consistent across independent datasets

There is an expectation that multiple, independent datasets that report molecular features across cancer patient cohorts with similar clinical profiles will show similar biological findings. If results are inconsistent between patient cohorts, differences in data processing and annotations, major batch effects or potentially biological effects could be the explanation. To test the consistency of our processed molecular measurements across patient cohorts, we evaluated patterns of transcriptome, copy number alterations, and mutations.

Gene expression, as measured by microarrays or RNA sequencing, is the most common molecular measurement in the *curatedPCaData* package (**Table 1**). To evaluate the consistency of expression patterns, we first performed a pairwise correlation analysis of gene expression differences in Gleason grade ≥8 vs. Gleason grade ≤6 tumor samples using the genes that were in common between the datasets (**Figure 1A**). Overall, we found that pairwise Pearson correlation between datasets was generally lowly correlated. Compared to the TCGA dataset^2^, the reported correlations were between 0.34 and 0.48 for Taylor et al.^4^, Weiner et al.^38^, Barwick et al.^39^, and IGC^40^. However, not all datasets were as correlated to TCGA. For example, the Friedrich et al.^41^ dataset only showed a correlation of 0.18, which could be attributed to difference in the underlying platform as gene expression in TCGA was measured by RNA sequencing and Friedrich et al. was measured by a custom Agilent microarray.

**Figure 1:**
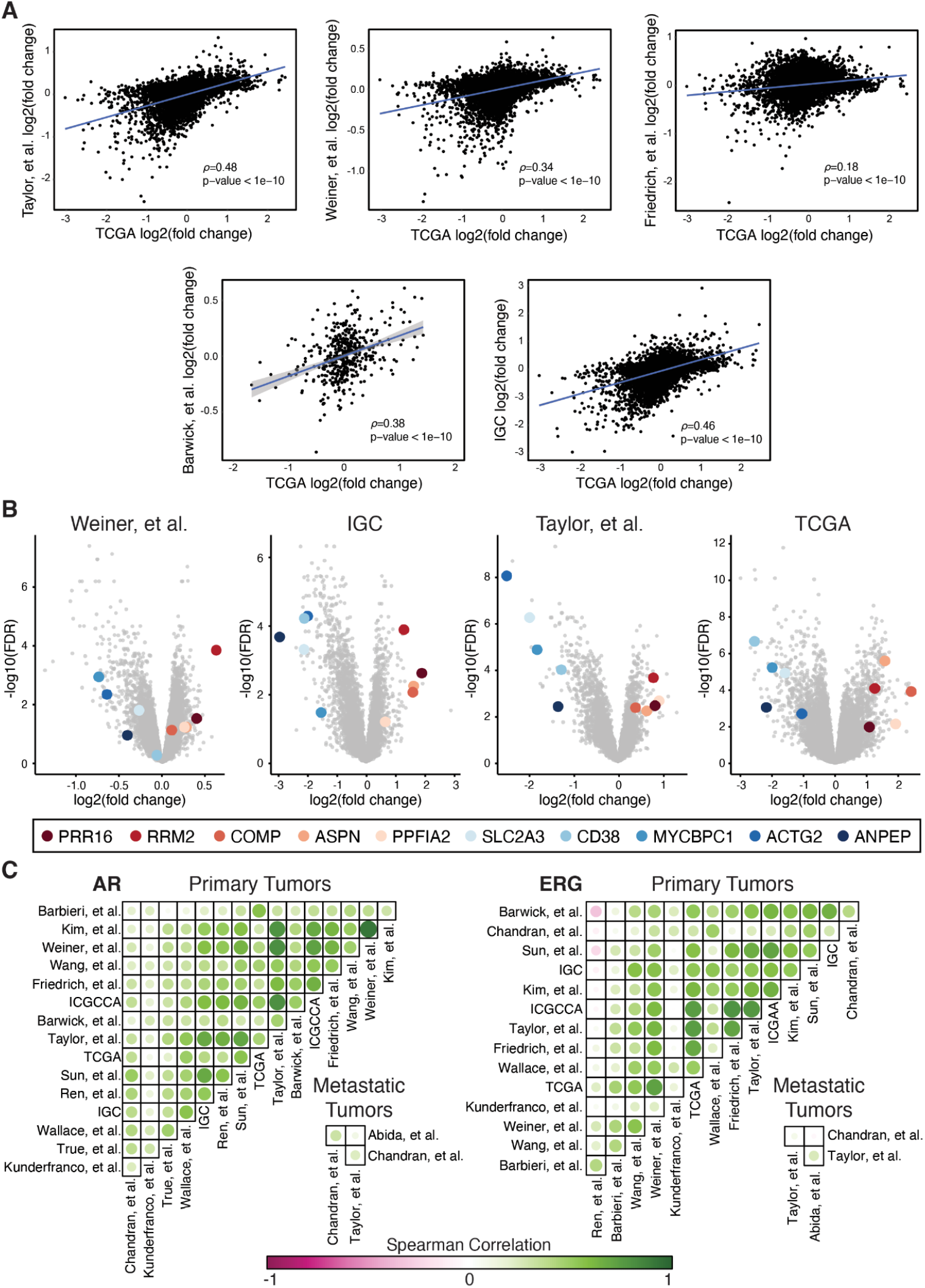
Gene expression patterns across datasets. **A**) Pearson correlation between datasets comparing differential expression of Gleason grade ≥8 vs. Gleason grade ≤6 samples for genes common between the datasets. **B**) Volcano plots for differential gene expression comparing Gleason grade ≥8 vs. Gleason grade ≤6 samples. The highlighted genes are the top five up- and down-regulated genes identified across the four datasets using Fisher’s method to combine p-values. **C**) Spearman’s rank correlations for all genes within the dataset were calculated compared to AR and the ETS transcription factor ERG. The Spearman correlation was calculated for the correlation patterns between datasets and displayed for AR (left side) and ERG (right side) in both primary and metastatic tumors.

Next, we identified the most commonly up- and down-regulated genes when comparing Gleason grade ≥8 vs. Gleason grade ≤6 tumor samples across multiple datasets (TCGA^2^, IGC^40^, Taylor et al.^4^, Weiner et al.^38^). We used the moderated t-test calculated through the *limma* R package to determine log fold change and p-values for individual datasets. We then integrated the four datasets using Fisher’s method to combine p-values to identify genes that were consistently up- (n=263) or down- (n=501) regulated and significant (q-value < 0.01) across these datasets (**Table S3**). Consistent with the biological processes associated with tumor growth and aggressiveness, the up-regulated genes are enriched for cell cycle-related processes, cell division, DNA replication, and DNA repair, while the down-regulated genes are enriched for positive regulation of apoptosis, negative regulation of ERK1 and ERK2 cascade, and cell-matrix adhesion. Using volcano plots for visualization, and for illustrative purposes, we highlighted the top 5 consistently up- (PRR16, RRM2, COMP, ASPN, PPFIA2) and top 5 consistently down-regulated genes (ANPEP, ACTG2, MYCBPC1, CD38, SLC2A3) (**Figure 1B**).

Finally, for gene expression, we evaluated the consistency of correlation patterns in relation to prostate cancer-associated genes. For each dataset, we calculated the Pearson correlation of all genes within the dataset to Androgen Receptor (AR) and the ETS transcription factor ERG. We then calculated the Pearson correlation of the correlation patterns to AR and ERG across datasets (**Figure 1C**). For the majority of datasets measuring gene expression in primary prostate tumors, the correlation patterns for AR across datasets were consistent with some datasets being highly correlated, such as Kim et al.^42^ and Weiner et al.^38^, or Taylor et al.^4^ and Sun et al^43^. Patterns for ERG expression were moderately to highly correlated, but there were some datasets with inverse correlation, such as Ren et al.^44^ and Sun et al.^43^, and Ren et al. and Barwick et al.^39^ While datasets with gene expression from metastatic tumors are few, the pattern of correlation between Chandran et al.^45^, Abida et al.^46^, and Taylor et al.^4^ were lower, likely due to the intrinsic heterogeneity of measuring gene expression from samples in the metastatic setting.

Prostate cancer is known to be heavily driven by copy number alterations which will impact the molecular measurements of gene expression. For datasets with copy number alteration information, *curatedPCaData* provides discretized copy number calls according to GISTIC2 (−2=deep loss, -1=shallow loss,0=diploid,1=gain, 2=amplification)^47^. We evaluated the overall copy number landscape and found that independent datasets showed highly similar patterns of copy number gain and loss in primary tumors (Taylor et al.^4^, TCGA^2^, Baca et al.^48^) (**Figure 2A**), with samples from metastatic tumors (Abida et al.^46^) showing an overall increase in copy number alterations as has been previously reported.^2,46^ We additionally evaluated the frequency of copy number alteration across several genes that have been shown to be associated with prostate cancer (PTEN, TP53, CHD1, MAP3K7, FOXA1, NXK3.1, USP10, SPOP^2,4,9,48–54^), along with the TMPRSS2:ERG fusion^2,55^. For these genes, we found the copy number alteration and mutation patterns to be consistent across datasets (**Figure 2B**, note that not all datasets have all genes measured for mutations or copy number). We also tested for patterns of co-occurrence and mutual exclusivity between these genes. While general patterns of co-alteration were consistent between datasets, the statistical significance, as measured in the primary tumor setting (Taylor et al.^4^, TCGA^2^, Baca et al.^48^), not surprisingly is highly dependent on the size of the dataset. In the metastatic setting (Abida et al.^46^), the frequency of alteration is consistently much higher and many genes are statistically significantly co-altered (**Figure 2B**).

**Figure 2:**
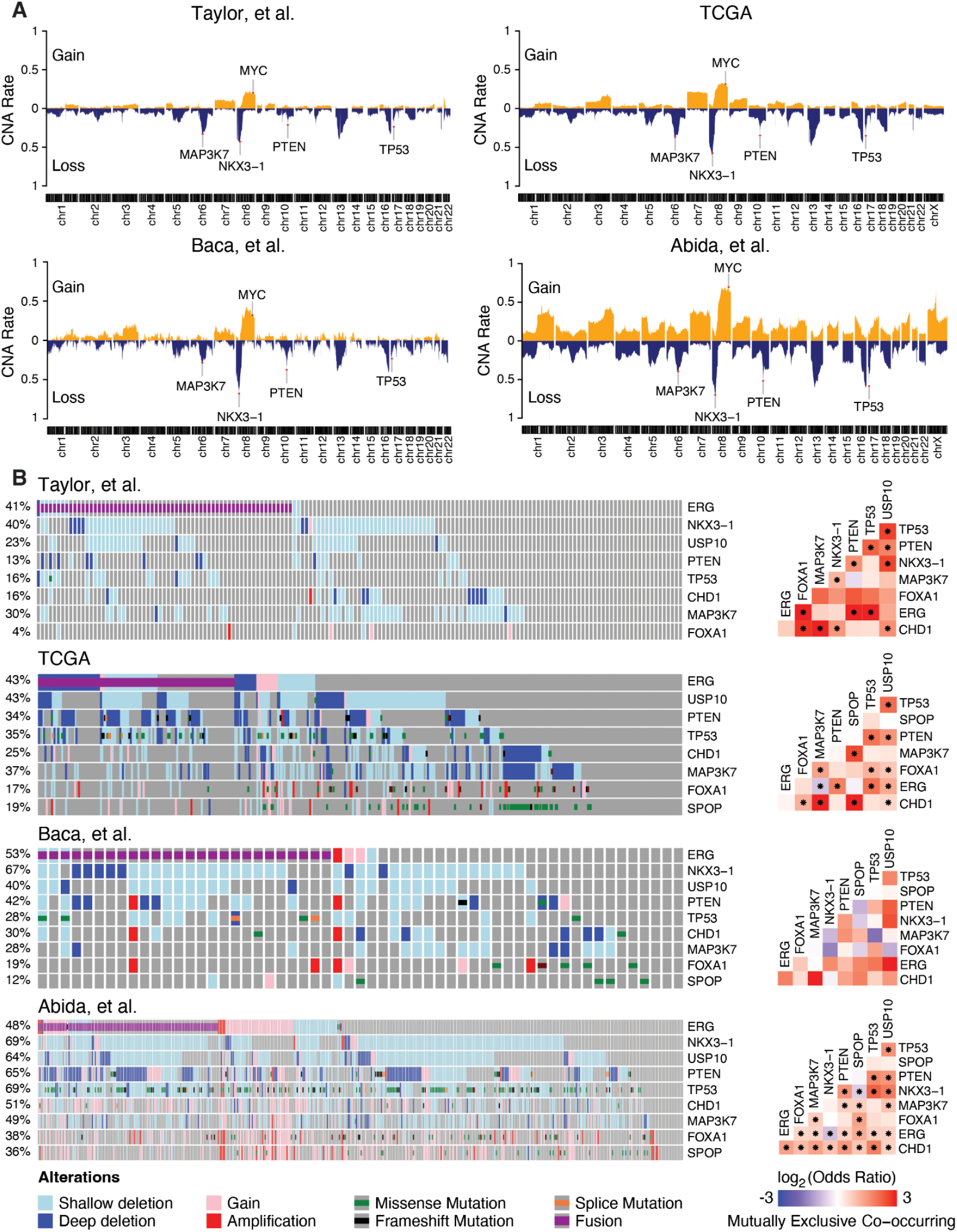
Copy number and mutational landscapes across datasets. **A**) Multiple known prostate cancer associated genes (MAP3K7, MYC, NKX3-1, PTEN, TP53) displayed consistent copy number loss/deletion or gain/amplification across datasets. **B**) Oncoprints (left side) for select prostate cancer associated genes are displayed across datasets. Mutual exclusivity (right side) was calculated using Fisher’s exact test (*p<0.05). Note that due to lack of overlap in omics, some alteration percentages combining CNA and mutations are under-estimated; for example Taylor et al.^4^ used a targeted sequencing panel and thus not all genes were measured for somatic mutations.

Overall, these benchmarking analyses show that the molecular features in primary prostate cancer are generally reliably and consistently measured across datasets. Gene expression patterns are correlated across datasets. Copy number results were more robust across datasets, with mutational information limited to a few datasets. The consistent data processing and harmonization of gene names across datasets provide a ready to use resource for meta-analysis.

### Derived features add value to published datasets

A value added in the *curatedPCaData* package, beyond data harmonization, is that features were systematically and consistently derived across datasets. Leveraging gene expression data, we inferred and evaluated estimates of risk (Oncotype DX^56^, Decipher^11^, and Prolaris^10^), AR scores, and microenvironment cell content leveraging the *Immunedeconv* R package^32^.

Prognostic risk scores are calculated from a select set of genes; thus missing genes and assay platform differences can impact the reliability of the computed scores^57^. To assess the impact of missing genes on risk score calculations, we benchmarked the risk scores included in *curatedPCaData* (Oncotype DX^56^, Decipher^11^, and Prolaris^10^) by removing different genes for calculating the risk scores, calculated the risk score with simulated missingness, followed by correlating the risk score derived from the incomplete gene set to the risk score calculated from the full gene list. Oncotype DX, a 12-gene signature, performed well overall when genes were missing from the gene list. As an example, with 5 genes missing over 100 random iterations, the average correlation coefficient was 0.891(median = 0.903) compared to the “ground truth” score using all genes (**Figure S2A**). Prolaris, a 34-gene signature, also proved to be highly robust whereby removing 10 random genes from the Prolaris gene list in the Kunderfranco et al. dataset had an average correlation with the original score of 0.973 (median = 0.974; **Figure S2B**). Decipher, a 17-gene signature, showed similar results to Oncotype DX where removing 5 genes resulted in an average correlation of 0.921 (median = 0.937; **Figure S2C**). Lastly, the AR score was calculated by taking the means across scaled gene expression values and found to be robust to the removal of genes. There are 20 genes that are used to calculate the AR score and we found that by removing 10 at random still provides an average AR score with a correlation of 0.930 (median = 0.935; **Figure S2D**).

In addition to prognostic risk and AR score calculations, we performed cell type deconvolution, which infers immune cells and other stromal cells from bulk tissue gene expression profiling. For datasets with gene expression, we calculated immune and other cell estimates using EPIC^27^, ESTIMATE^28^, MCP-counter^29^, quanTIseq^30^, and xCell^31^ as implemented in the *immunedeconv* R package^32^, and CIBERSORTx^34^. While deconvolution methods vary in the types of cells that they estimate, the overall methodology has been shown to produce robust predictions and comparison between methods have been shown to be mostly consistent and robust, which is covered in depth by Sturm et al.^32^ and was a major motivation to develop the *immunedeconv* R package. The following section highlights how the inferred cell content can be used to infer associations with clinical outcomes using *curatedPCaData*.

### Endothelial cell content predicts patient outcomes

Leveraging the results from the immune and cell deconvolution methods from bulk transcriptome data, we evaluated the relationship between inferred cell types, patient outcomes, and disease progression. We found that the estimates of endothelial cell content as estimated by xCell^31^, MCP Counter^29^, and EPIC^27^ were predictive of biochemical recurrence. It was encouraging to also find that the results from the three independent methods were highly correlated (**Figure 3A**), which provides support that the signal is reproducible and not an artifact of one deconvolution method. For illustrative purposes, we stratified patients in the TCGA^2^ and Taylor et al.^4^ cohorts into the top ⅓ and bottom ⅔ by endothelial cell estimates. The endothelial cell scores were dichotomized at the upper tertile, and HRs were estimated using univariate Cox models for each method (EPIC, MCP-counter, and xCell) by comparing upper tertile with the two lower tertiles in order to make sure that the binarized endothelial cell score statuses were comparable between methods. We noted that the univariate Cox models agreed on the Hazard Ratio (HR) estimates and statistical significance across the methods and datasets, with HR estimates ranging between 2.02 to 2.446 in TCGA and 1.959 to 3.536 in Taylor et al. (**Figure 3B**). When Gleason grade group (≤6, 7, ≥8) was modeled as a univariate Cox model predictor, its unit increase estimate for HR was of similar effect size as having the top tertile for endothelial cells with 2.154 and 3.52 for TCGA and Taylor et al., respectively. Patient samples with a high endothelial score show significantly shorter times to biochemical relapse (**Figure 3C**). Furthermore, we evaluated primary tumor datasets for the association between endothelial cell estimates and Gleason grade. Across the datasets that reported at least 10 patients per Gleason grade group and where we could infer endothelial cell content from gene expression data (TCGA^2^, Taylor et al.^4^, Friedrich et al.^41^), we consistently found increased estimated presence of endothelial cells in Gleason grade ≥8 compared to Gleason grade 7 or ≤6 (**Figure 3D**).

**Figure 3:**
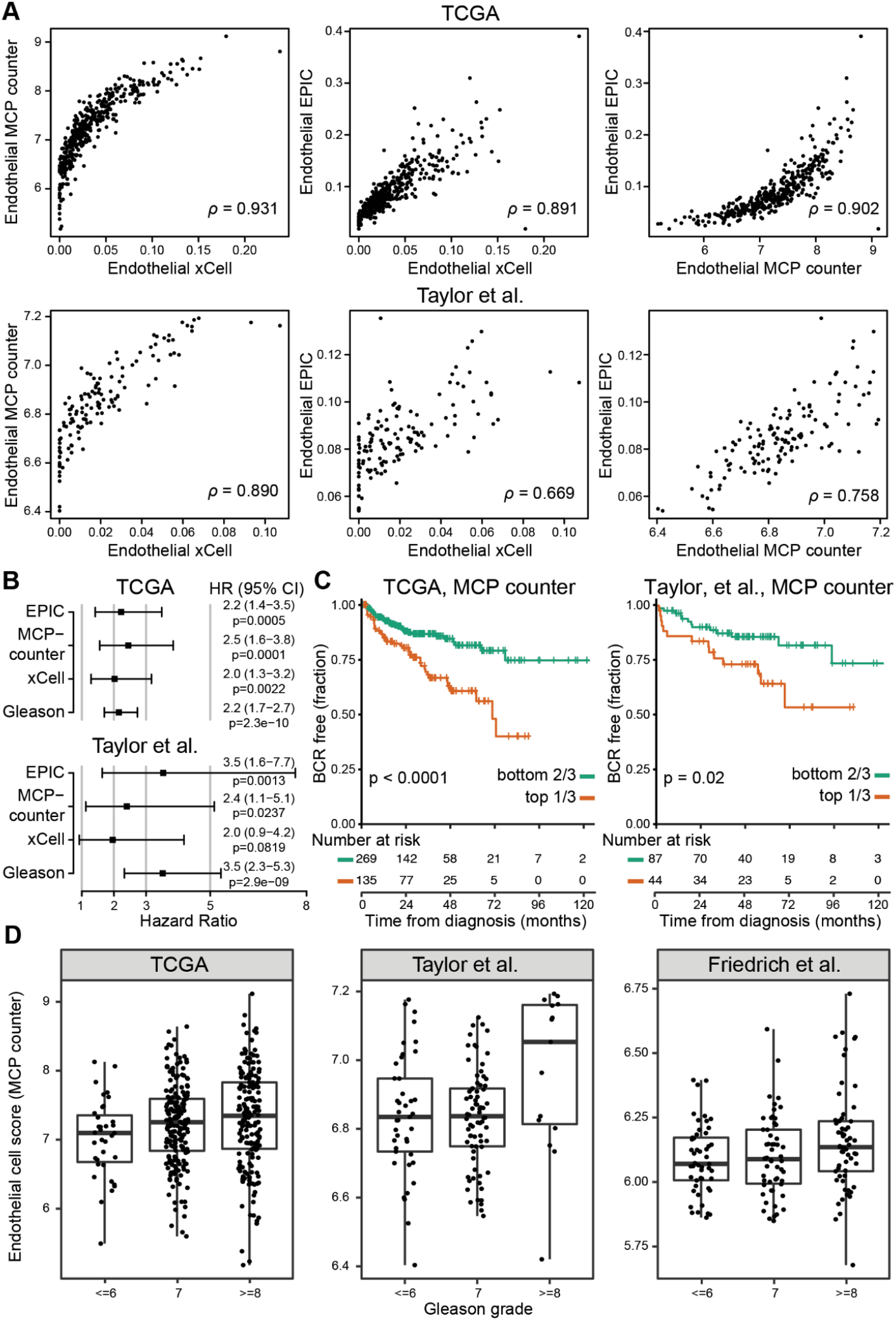
Estimates of endothelial cell content are associated with clinical outcomes. **A**) The endothelial cell scores calculated from gene expression across TCGA^2^ and Taylor et al.^4^ were highly correlated (Spearman correlation) across the three estimation methods, EPIC^27^, MCP-counter^29^, and xCell^31^. **B**) Forest plots for univariate Cox proportional hazard models illustrate that all three methods and Gleason grade were predictive of biochemical recurrence. **C**) Endothelial cell score top tertiles, as illustrated using MCP-counter’s estimates, showed a statistically significant stratification for worse outcome in TCGA and Taylor et al. datasets. **D**) In addition to being associated with biochemical recurrence, the estimates from MCP-counter are associated with tumor Gleason grade groups.

It has been established that the cellular content of the tumor microenvironment can be predictive of tumor progression and response to treatment, mostly in the context of immune cells^33^. Similarly, angiogenesis and the vascularization of the tumor microenvironment has been associated with tumor progression and outcomes^58–61^, with specific studies linking endothelial cell content to prostate cancer aggressiveness^62,63^. Our findings are consistent with previous results and demonstrate the strength of leveraging the inferred features across multiple, independent datasets through *curatedPCaData*.

## DISCUSSION

The *curatedPCaData* R package provides a harmonized and centralized resource for prostate cancer studies with multi-omic and clinical data that can be leveraged easily for cancer research. The cross study analyses presented herein demonstrate the strength of leveraging multiple studies in prostate cancer; however, it is important to understand and incorporate relative differences between studies, their aims, design and the underlying composition in such data analysis. For example, Abida et al.^46^ focused on the progressed metastatic form of the disease and reported a significant number of disease related deaths suitable for death-related survival modeling. On the other hand, Friedrich et al.^41^, Hieronymus et al.^6^, ICGC-CA^64^, and TCGA^2^ also reported overall survival, but they present a more indolent form of the disease with a lower count of deaths, making survival modeling more challenging. Furthermore, biochemical recurrence is often used as a surrogate for progression free survival and is reported in Barwick et al.^39^, Sun et al.^43^, Taylor et al.^4^ and TCGA^2^; of these four datasets we focused our Cox models for recurrence on Taylor et al. and TCGA, as Barwick et al. used a very targeted custom DASL gene panel (<1,000 genes) making cell composition estimation unreliable for most methods. Sun et al. only report recurrence as a binary outcome without follow-up times, rendering it not suitable for Cox proportional hazards models or survival estimation using Kaplan-Meier method. Despite the differences in reported variables, a considerable amount of clinical information is made available across independent datasets to draw associations with molecular features.

Researchers should also consider the original study aims, as these will be reflected in which metadata fields and omics that will be available. For example, Weiner et al.^38^ studied ethnicity related PCa-trends, thus the patients had accurate demographics-related metadata commonly available, while samples were just described as being primary tumors. In contrast, Wang et al.^65^ studied how sample composition (tumor cells, stroma, atrophic grand, or benign prostate hyperplasia) could be differentiated based on gene expression, thus providing metadata suitable for tumor purity estimation, but provided no clinical end-points or patient characteristics. While we have gone through great effort to minimize technical and reporting variability, some fundamental study characteristics will inevitably be not comparable. Thus, combining studies ought to be planned with care to avoid introducing confounding effects. To this end, *curatedPCaData* offers assistance in bringing together studies suitable for efficiently tackling specific prostate cancer related research questions.

Additional consideration should be given to how studies reported the common end-point of Gleason grade. In *curatedPCaData*, we provided summarized results across studies as Gleason grade groups (≤6, 7, ≥8), though studies might have additional information to report. For example, Weiner et al.^38^ reported an International Society of Urologic Pathologists (ISUP) disease stage ranging from 1-5, for which the suggested mapping to the traditional Gleason grade was done^66^. Multiple studies reported Gleason as the sum of major + minor Gleason grades or a grade group (≤6, 7, ≥8), thus groupings were offered as an endpoint with equal level of granularity, while finer level of detail was offered in alternate clinical metadata columns when available. In ambiguous cases, the primary publications and the supplementary material was mined, along with contacting the primary authors in many cases, in an effort to offer accurate and up-to-date information on both the clinical metadata and the primary data. For this purpose, a great deal of manual labor was required to curate the *curatedPCaData* datasets. The resulting datasets were thus standardized to be as comparable as possible, while retaining details essential to the studies. To this end, we offer a great variety of R package vignettes alongside *curatedPCaData* with numerous examples and extra data characteristics, which assist the end-user in planning their analyses (**Table S2**).

One benefit of the *curatedPCaData* is that it greatly lowers the barrier for accessing data to rapidly test hypotheses and generate novel hypotheses supported by multiple, independent datasets. The code used to generate the MAE objects is offered within the R package and GitHub repository as supplementing code. The processed MAE objects exported from the package are the main focus of the package; however, from a developer point of view they also offer natural potential for future extensions such as: a) adding new studies and exporting them as new MAE objects using the pipelines developed in *curatedPCaData*; b) supplementing the existing MAE slots with newly derived variables or even adding other primary omics data; c) extending the existing clinical metadata fields to include new fields.

Currently, *curatedPCaData* offers a base R Shiny^67^ interface to the package as well, with plans to extend the visual browser-based access to the data. While on-going efforts such as the NCI Genomic Data Commons^68^, cBioPortal^69^, or the International Cancer Genome Consortium^70^ already aim at providing a standardized approach to tackling complex omics traits in cancer, *curatedPCaData* is the first harmonized, multi-study, hands-on data resource intended for analysts with a strong focus on prostate cancer and allowing for maximum flexibility of the analyses, using the R statistical software^71^. As such, the presented proof-of-concept analyses provide merely a staging platform for more efficient exploration of multi-omics signatures coupled with clinical metadata for the wider research community for prostate cancer.

## METHODS

### Data acquisition

Gene expression, copy number alterations and mutation data were downloaded from Gene Expression Omnibus (GEO)^72^ using *GEOquery* (R package version 2.64.2) and from cBioPortal^69^ using *cBioPortalData* (R package version v2.8.2) and *cgdsr* (R package version v1.3.0) (**Figure S1A**). In addition to downloading raw data from GEO, *GEOquery* was used for downloading the latest array-specific annotations and all three R packages were further utilized to download clinical metadata accompanying the raw data. Raw CEL-file files for Affymetrix-arrays were RMA-normalized in *oligo* (R package version v1.62.1) with functions *read*.*celfiles, rma, getNetAffx*, and *exprs*. Agilent arrays were processed using *limma* (R package version v3.52.2) with the functions *read*.*maimages, backgroundCorrect, normalizeBetweenArrays*, and *avereps*. For custom arrays such as the DASL array in Barwick et al.^39^, quantile normalization was used together with log-transformation. No additional normalization was done on the gene expression data from cBioPortal, since cBioPortal offers pre-normalized data. For data with raw copy number alteration available, these were processed using *rCGH* (R package version v1.26.0) with functions *readAgilent, adjustSignal, segmentCGH*, and *EMnormalize*. This yielded log-ratios, which were input to GISTIC2^47^ when available. Copy number alteration matrices from cBioPortal with pre-existing GISTIC2 calls were stored with the discretized calls consistently across all the datasets.

The TCGA Prostate Cancer (PRAD) dataset was downloaded from Xena Browser^73^, due to better data quality and providing tumor samples and normals separately, instead of providing relative tumor to normal gene expression found in cBioPortal processed data. We also removed low-quality samples which were excluded from the TCGA publication due to RNA degradation from the gene expression matrix to provide users with the most reliable information. We followed uniform naming conventions for all the metadata fields and leveraged data in the original publications to obtain maximum information in case information wasn’t readily available in these public repositories (**Table S1**).

All layers of data, namely the gene expression, copy number alterations and mutations, underwent a harmonization process to ensure uniform gene naming conventions. Note that some datasets have matched normal samples to call somatic mutations and some datasets do not have matched normal samples and are thus tumor-only variants. The mutation calling status is noted in the “Mutation_status” field.The latest hg38 gene symbols, aliases and locations were downloaded using *biomaRt* (R package version v2.52.0). We then mapped all the gene names to the up-to-date dictionary to ensure consistency in HGNC symbols across all datasets. A liftover from hg19 to hg38 was done as part of the harmonization using the liftOver function from *rtracklayer* (R package version v1.56.1), for mutations called with an older genome assembly to ensure uniformity.

Clinicopathological features were processed using R scripts customized to each dataset. Features were collected from supplementary annotation files and processed to map features to the data dictionary (**Table S1**). The data dictionary ensured common terminology and some additional features, such as Gleason grade group (where not supplied by the primary publication), were inferred using a predefined set of rules. The scripts for each dataset are made available in *curatedPCaData*.

### Derived features

A number of derived features were computed for the final MAE-objects (**Figure S1B**). Using gene expression data, we calculated cell proportions, genomic risk scores, and AR scores. The *immunedeconv*^*32*^ (R package version v2.1.0) wrapper package was used to estimate cell proportions from EPIC^27^, ESTIMATE^28^, MCP-counter^29^, quanTIseq^30^, and xCell^31^. As the implementation of CIBERSORTx^34^ required external access using the free academic license, it was run with default parameters on their web interface and quantile normalization disabled with the normalized gene expression data as input and LM22 signature matrix used to infer cell types. The output CIBERSORTx matrices were then downloaded and integrated into the MAEs.

Due to the different platforms (sequencing, different brands and versions of microarrays) used to assess gene expression, not all datasets have the same set of genes. To determine the impact of gene missingness on the precomputed scores that this would have on those studies without all genes, we benchmarked the Oncotype DX^56^, Decipher^11^, and Prolaris^10^ risk scores and the AR score. This was performed by identifying the study in *curatedPCaData* that contained the most genes belonging to the score. By using this study we were able to get as close to what the true score value would be. Assessing the impact of missing genes was performed by randomly removing genes to simulate missing between 1 and 10 genes for Prolaris^10^ risk score (34 genes in the complete signature) and AR score (20 genes), and removing between 1 and 5 for Oncotype DX^56^ and Decipher^11^ risk scores (12 and 20 genes, respectively). Since the number of gene combinations that can be made by simulating 10 missing genes for a risk score such as Prolaris^10^ is large, the combinations were sampled to cut down on vignette and package build time. The number of combinations used for assessing impact of missingness in Decipher^11^, Oncotype DX^56^, and AR scores was 100 while Prolaris risk score used 50 combinations.

We implemented the Oncotype DX^56^, Decipher^11^, and Prolaris^10^ risk scores based on the instructions in their original publications supported by the implementation outlined in Creed et al.^57^ The gene list (n=12 matching genes) for Oncotype DX matched perfectly with several studies: Abida et al.^46^, Kim et al.^42^, Ren et al.^44^, Sun et al.^43^, Taylor et al.^4^, TCGA^2^, Wallace et al.^74^, and Weiner et al.^38^ We considered TCGA to be the most complete dataset as well as most widely used, thus we used the gene expression from TCGA for testing the variability of the Oncotype DX score due to missing genes (**Table S4**). The gene list (n=17 matching genes) for Decipher did not have a 1-to-1 match with any study in *curatedPCaData*, but did have the highest number of matching genes in Ren et al.^44^ (18 genes were a 1-to-1 match with two genes from Decipher missing) while Abida et al.^46^, Friedrich et al.^41^, and TCGA^2^ had slightly fewer number of matching genes (17 genes were a 1-to-1 with 3 genes missing). We used TCGA gene expression for benchmarking inferred risk scores from Decipher. Prolaris required the largest number of genes (n=34 matching genes) to calculate risk. Kunderfranco et al.^75^ had the highest number of matching genes with 32 1-to-1 matches and only 2 genes missing. The next highest 1-to-1 match was ICGC^64^ where 29 genes were 1-to-1 matches. Because of the high number of matching genes, we selected Kunderfranco et al. as the benchmarking study for Prolaris (**Table S4**).

AR-scores were calculated for the 20 genes identified originally in Hieronymus et al.^76^ and then calculated as the sum of z-scores of AR signaling genes as described by TCGA^2^. There were 8 studies that matched all 20 genes used to calculate the AR score; we leveraged TCGA gene expression for benchmarking.

### Statistical analysis

While the primary focus is on providing readily processed MAE-objects with *MultiAssayExperiment* (R package version v1.21.6), *curatedPCaData* delivers several application examples as R vignettes and documentation, with relevant statistical methodology applied there-in (**Table S2**). Cox proportional hazard models and Kaplan-Meier (KM) curves were fitted with *survival* (R package version v3.3-1) and plotted using *survminer* (R package version v0.4.9), and the corresponding p-values were calculated using log-rank tests.

Differential gene expression was calculated as the average log-transformed expression of Gleason grade ≥8 samples minus the average log-transformed expression of Gleason grade ≤6 samples. Statistical significance was determined by comparing the log-transformed gene expression of Gleason grade ≥8 compared to Gleason grade ≤6 samples using the moderated t-test as implemented in *limma* (R package version v3.52.2). The final p-values were adjusted for multiple testing using the Benjamini & Hochberg correction. Pearson correlation was used to compare differential expression in **Figure 1A**. The genes reported in **Figure 1B** were identified using Fisher’s method to combine p-values for statistical significance. The log fold change was then tested to ensure consistent up- and down-regulation of the associated gene, meaning a gene needed to have logFC > 0 or logFC < 0 across all four datasets tested. The top up- and down-regulated gene sets were tested for pathway and biological process enrichment using the DAVID web server^77^. The correlations reported in **Figure 1C** were calculated using Spearman’s rank correlation.

Genes were defined to be co-occurring or mutually exclusive based on the odds ratio (OR) which is calculated as: OR = (Both* Neither) / (B Not A * A not B) where A and B stand for alterations in A and B respectively. We define any alteration in copy number or mutations that are not silent as an alteration. The significance of mutual exclusivity/co-occurrence was computed using the Fisher’s Hypergeometric Test and the Benjamini-Hoschberg correction was applied to determine the adjusted p-values. Mutual exclusivity plots for different data sets shown in **Figure 2B** (right side), provide information on whether or not a set of important genes in PCa are significantly altered together.

Statistical modeling used to identify interesting derived features predictive of biochemical recurrence were based on 10-fold cross-validation (CV) of Cox models regularized using LASSO using *glmnet* (R package version v4.1-4)^78^. There were three methods that calculated endothelial cell abundance scores (EPIC^27^, MCP-counter^29^, and xCell^31^). Among these methods, endothelial cell abundance scores were predictive in at least one of these datasets, when predictive features were chosen according to the optimal regularization coefficient λ in the CV-curve.

Spearman’s rank correlation was used to assess the non-linear association between endothelial cell scores in **Figure 3A**. Cox proportional hazards models were fit as univariate models with biochemical recurrence as an endpoint, by introducing one of the endothelial scores at a time to a separate model, compared with using Gleason score sum as an univariate predictor; these were then plotted together as a forest plot in **Figure 3B**.

## DATA AVAILABILITY

All the data presented here-in are available as MultiAssayExperiments^1^ in the *curatedPCaData* R package, along with code that can be used to reproduce these objects. The original raw data repositories along with unique identifiers are listed, such as GEO accession ids or cBioPortal identifiers listed in **Table 1**.

## CODE AVAILABILITY

All the code used to generate the processed datasets, as well as the resulting R package are available openly at: https://github.com/Syksy/curatedPCaData

## Acknowledgements

This work is supported by grants CA242747 to J.C.C., S.T., and B.F., CA231978 to J.C.C., the Finnish Cultural Foundation and the Finnish Cancer Institute as FICAN Cancer Researcher to T.D.L., in part by the Biostatistics and Bioinformatics Shared Resource at the H. Lee Moffitt Cancer Center & Research Institute, an NCI designated Comprehensive Cancer Center (P30CA076292), and in part by the Biostatistics and Bioinformatics Shared Resource at the University of Colorado Cancer Center, an NCI designated Comprehensive Cancer Center (P30CA046934). The authors would like to extend gratitude to the curated datasets’ original authors, who provided irreplaceable advice and additional information for their studies.

## AUTHOR CONTRIBUTIONS

T.D.L., V.S., A.S., J.C., F.C.F.C., K.S., C.C.L. developed and wrote the R package, documentation and constructed the exported data objects; T.D.L., V.S., J.C., F.C.F.C., C.C.L., T.G., S.T., J.C.C. designed the harmonized data processing pipeline; T.D.L., V.S., A.S., M.O., B.F., S.T., J.C.C. contributed R vignettes; T.D.L., V.S., J.C., A.S., J.C., F.C.F.C., K.S., T.G., B.F., S.T., J.C.C. contributed original analyses; T.D.L., V.S., A.S., M.O. visualized data and analyses; T.G., B.F., S.T, J.C.C. supervised the project and obtained funding; T.D.L., V.S., A.S., S.T., J.C.C. drafted the manuscript; All authors read and approved the final manuscript.

## COMPETING INTERESTS

The authors declare the following competing interests: J.C.C. is co-founder of PrecisionProfile and OncoRX Insights. All other authors declare no competing interests.

## SUPPLEMENTARY MATERIAL

### Supplementary Tables

**Table S1**: Template used for extracting data for the PCa clinical metadata colData-slots in each MAE-object. Also exported from the package namespace via *curatedPCaData::template_prad*.

**Table S2**: Vignettes provided alongside *curatedPCaData* (≥ v1.0), topics and aims

**Table S3:** Differential expression of the four datasets (TCGA^2^, IGC^40^, Taylor et al.^4^, Weiner et al.^38^) in **Figure 1B** with the genes that are commonly and significantly up- and down-regulated identified.

**Table S4:**The intersection between Prolaris, Oncotype DX, Decipher, and Androgen Receptor (AR) score’ genes and genes that are found in studies within curatedPCaData R Package. A gene from the score or its aliases matched either with a single gene in the dataset (1-to-1 match), gene from the score matched or its aliases had multiple matches in the dataset (1-to-many), or the gene from the score calculation was missing from the dataset altogether.

## Supplementary Figures

**Figure S1:**
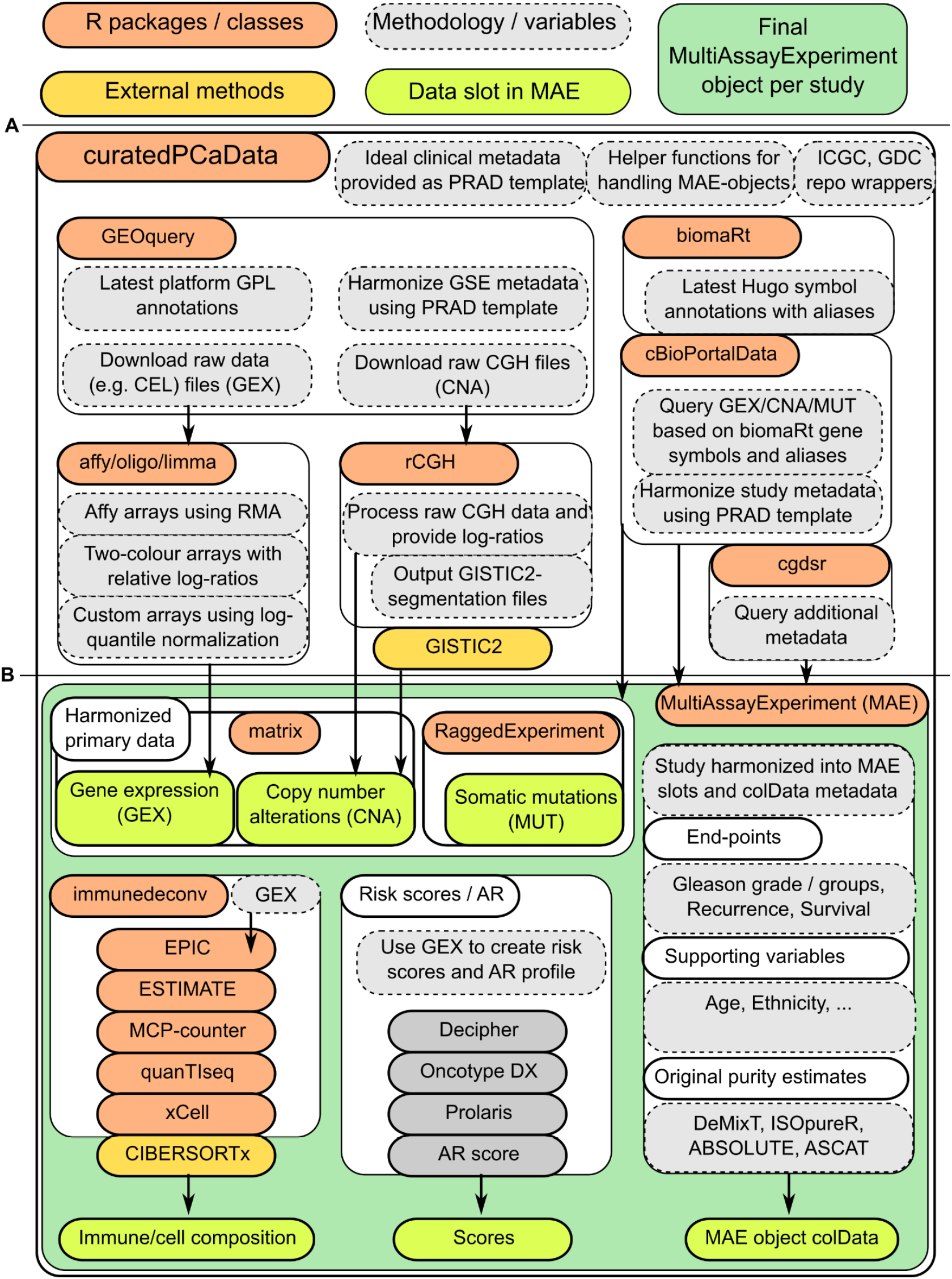
Workflow of the *curatedPCaData* MultiAssayExperiment-object generation. **A**) Primary raw data is extracted mainly using the GEOquery and cBioPortalData packages. Raw data are processed according to latest annotations with the help of biomaRt and assay-specific packages, and then processed using affy, oligo, limma, and rCGH packages where appropriate; **B**) MAE-object is constructed while providing access to the primary data (GEX, CNA, and MUT), offering derived variables (decompositions and scores), and corresponding clinical metadata (MAE colData-slot)

**Figure S2:**
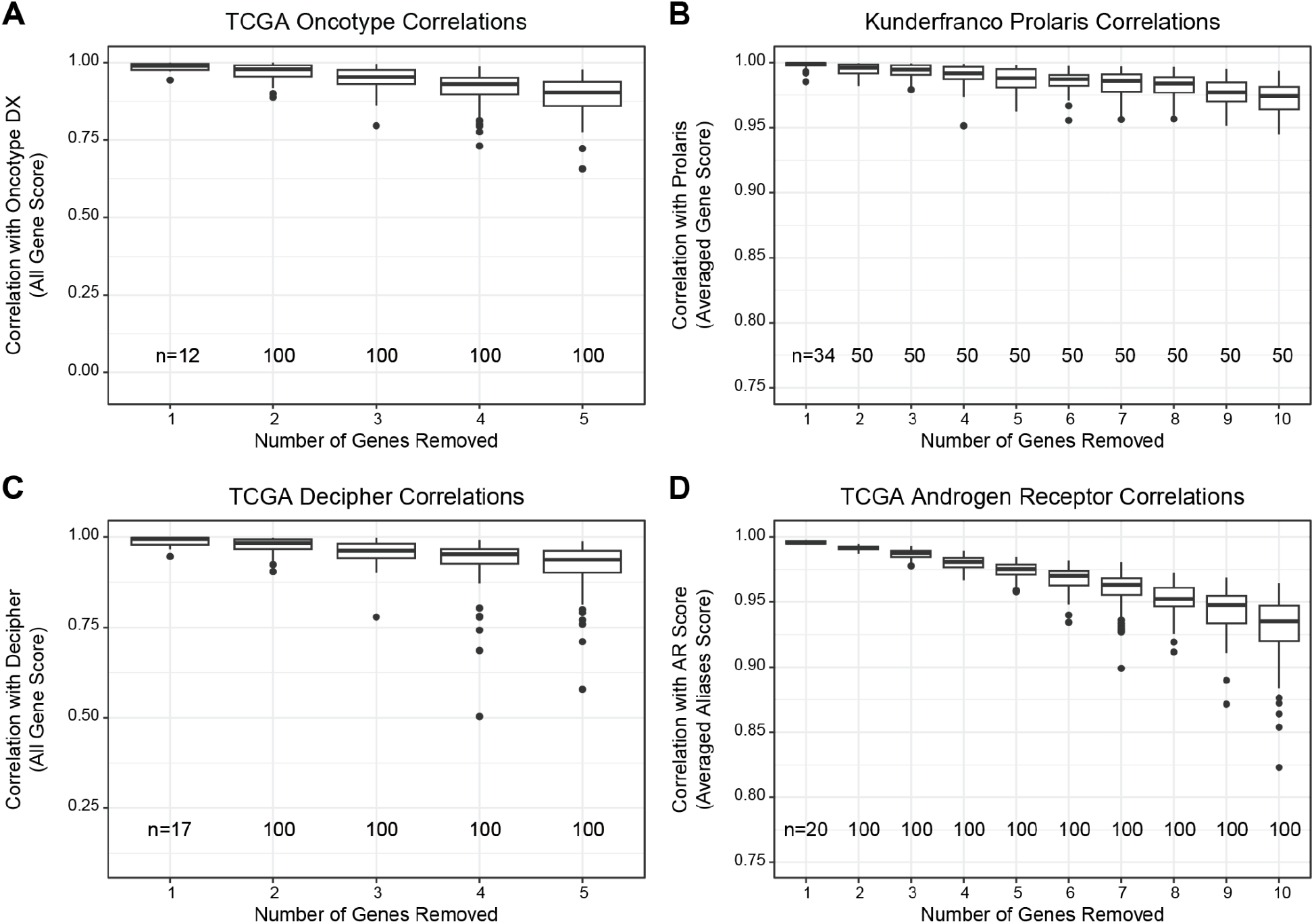
Impact of Gene Missingness on Risk and AR Score Reliability. Prostate risk scores and AR score were benchmarked using datasets from the *curatedPCaData* package to determine how missing genes impacted their reliability. The number of trials are listed at the bottom of each figure panel. **A**) TCGA was used to assess Oncotype DX risk score removing between 1 and 5 genes. **B**) Kunderfranco et al. was used to assess Prolaris risk score by removing between 1 and 10 genes. TCGA was leveraged to assess gene removal for **C**) Decipher (1-5 genes) and **D**) Androgen Receptor (1-10 genes).

## Notes

https://github.com/Syksy/curatedPCaData

